# The Investigation of Somatostatin Receptors as a Potential Target in Breast Phyllodes Tumours

**DOI:** 10.1101/2024.01.21.576520

**Authors:** Hande Süer Mickler, Murat Mert Erkan

## Abstract

Somatostatin receptors (SSTRs) constitute a family of G protein-coupled receptors that modulate hormonal secretion and regulate cell proliferation and apoptosis upon binding with somatostatin or its synthetic analogues. SSTRs are expressed in most neuroendocrine neoplasms, particularly in gastroenteropancreatic neuroendocrine tumours, and have been utilized as diagnostic markers as well as therapeutic targets. The radioiodinated somatostatin analogue 1,4,7,10-tetraazacyclododecane-1,4,7,10-tetraacetic acid-Tyr3-octreotate (DOTATATE) has been employed for SSTR-targeting for either diagnostic or therapeutic purposes depending on the labelling with ^68^Gallium or ^177^Lutetium, respectively.

SSTR expression is reported in a subset of breast adenocarcinoma and breast neuroendocrine carcinomas; however, minimal knowledge exists regarding their expression in fibroepithelial (biphasic) breast lesions such as fibroadenoma and phyllodes tumours. Aggressive ends of the spectrum, i.e. “cystosarcoma phyllodes” may present a management challenge with recurrences and metastases. In this exploratory study, we investigated both gene and protein expression of SSTR in fibroepithelial lesions of the breast. Our findings reveal that both fibroadenoma and phyllodes tumours express SSTRs. Immunohistochemical analyses suggested that this expression is in the stromal, not epithelial, component by demonstrating that SSTR is predominantly expressed in the areas overlapping with α-smooth muscle actin-positive myoepithelial cells around blood vessels and capillary structures. This study is the first in the literature to demonstrate SSTR positivity in mammary fibroepithelial neoplasms., In a patient undergoing a gallium scan, which is widely used in clinical practice, if a positive lesion is discovered in the breast, the possibility of a fibroadenoma or phyllodes tumour needs to be considered. Once validated, these findings may also have significant implications for the management of these tumours.

## INTRODUCTION

Somatostatin is a widely distributed peptide in both the central and peripheral nervous system and tissues such as the pancreas, gut, adrenals, thyroid, kidney, and immune system, regulating a variety of physiological functions^1, 2^. While it acts as a neurotransmitter in the nervous system, its hormonal activities include inhibition of the secretion of various hormones, including growth hormone, insulin, glucagon, gastrin, serotonin and calcitonin^3^.

The biological effects of somatostatin are mediated by high-affinity plasma membrane receptors known as somatostatin receptors (SSTR)^1, 2^. There are five subtypes of human SSTRs with a similar affinity to bind naturally occurring somatostatin 14 and 28. Due to the short half-life of natural somatostatin (2-3 minutes), many somatostatin analogues with longer half-lives have been synthesised^1^. Among them, long-acting release forms; octreotide, lanreotide and vapreotide are widely used in the clinic^1^. These synthetic analogues have different affinities for different SSTRs; for example, octreotide has a high affinity to SSTR2, moderate affinity to 3 and 5, and lower affinity to SSTR1 and SSTR4^3^. Among these SSTR1 and SSTR2 were found to be consistently expressed in neuroendocrine neoplasms of the gastrointestinal/ pancreatobiliary tract, and this discovery has led to revolutionary changes in the diagnosis and management of these tumours^4-6^. In addition to its therapeutic usage as a competitive inhibitor of somatostatin at the receptor level, octreotide is frequently used for SSTR imaging of neuroendocrine tumours. ^68^Gallium (^68^Ga)-labelled 1,4,7,10-tetraazacyclododecane-1,4,7,10-tetraacetic acid-Tyr3-octreotate (DOTATATE) has been used as a radiotracer for diagnostic imaging of SSTR overexpressing tumours, particularly gastrointestinal- and pancreatic-neuroendocrine tumours^7^. DOTATATE uses octreotide to target SSTR, while ^68^Ga, a generator-produced positron emitter that allows on-site, on-demand availability, provides a tracer signal when imaging with positron emission tomography/ computed tomography (PET/CT)^7^. DOTATATE backbone is also used in theranostics for radionuclide therapy when chelated with ^177^Lutetium (^177^Lu)^7^.

The incidental detection of breast cancer by ^68^Ga-DOTATATE has been observed in a few studies over the last decade. Up to 50% of breast adenocarcinoma and primary or metastatic neuroendocrine carcinomas of the breast demonstrated increased uptake of ^68^Ga-DOTATATE; however, there is very little information on the expression of SSTR in fibroepithelial lesions of the breast^8, 9^. Papadakis et al. observed a positivity of ^68^Ga-DOTATATE in the breast, later confirmed to be a fibroadenoma while scanning for metastatic neuroendocrine tumours^10^. Another study retrospectively evaluated 1573 patients for incidental uptakes of DOTA-PET and found that 6 out of 12 patients were considered fibroadenoma, with 4 of them clinically proven by biopsy^11^.

Fibroepithelial lesions are biphasic tumours of the breast, containing stromal and epithelial components and showing differences in biological behaviours and clinical management. They include common benign fibroadenomas and rare phyllodes tumours with a spectrum of locally aggressive neoplasia to high-grade sarcomas (also called “cystosarcoma phyllodes”) with a risk of metastasis and death^12^. Even though the diagnosis of fibroadenoma is straightforward, it can be challenging for phyllodes due to overlapping histological features; hence, they can be mistaken for fibroadenomas^12^. Clinical management also differs greatly, while conservative treatments with follow-up are recommended for most fibroadenomas; phyllodes tumours are generally resected surgically^12, 13^. Since phyllodes tumours have a potential for recurrence and progression, curative therapy requires tumour-free margins^13^. Depending on the size and localisation of the lesion, to achieve tumour-free resection, a complete mastectomy might be necessary^13^.

We also observed an incidental breast mass with ^68^Ga-DOTATATE uptake in one of our patients who was diagnosed with metastatic pancreatic neuroendocrine tumours (PanNET) (Figure 1). As a part of the treatment regimen, this patient received adjuvant ^177^Lu-DOTATATE treatment after pancreaticoduodenectomy, lymphadenectomy, and right hepatectomy. Importantly, during ^177^Lu-DOTATATE treatment, the size of the breast mass also decreased over time (Figure 1A-E)

**Figure 1:**
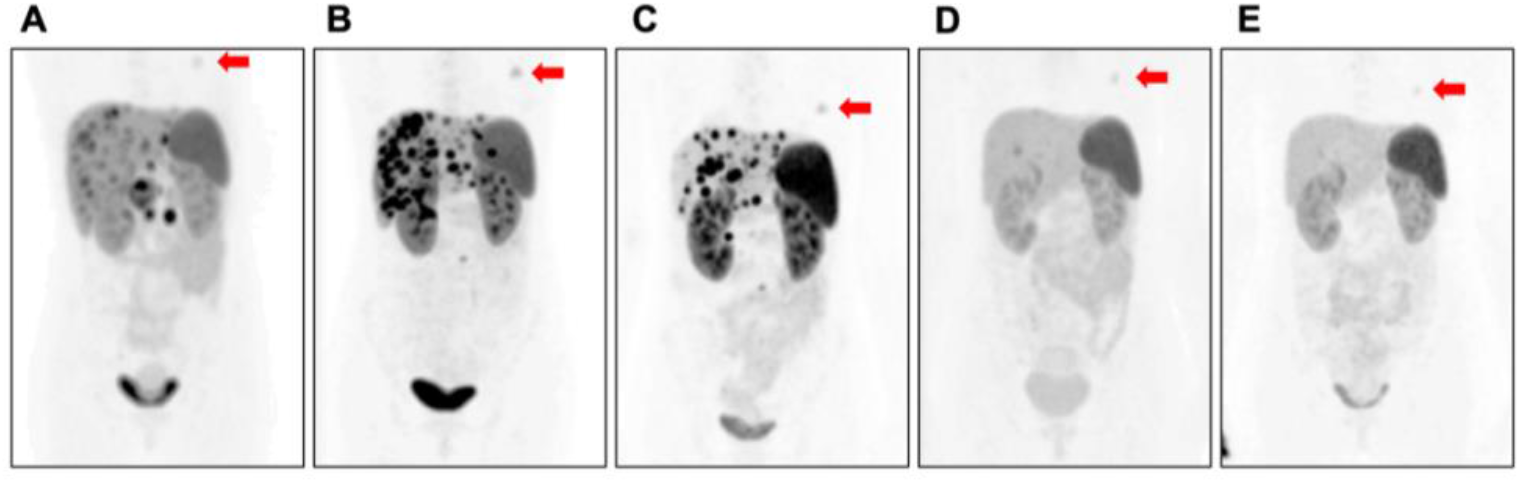
A. Pre-operative B. Post-pancreaticoduodenectomy and lymphadenectomy C. Post-right hepatectomy D. 4^th^ 177Lu-DOTATATE Treatment E. 5^th^ 177Lu-DOTATATE Treatment. Note the gradual decrease in the Ga-68 uptake of the breast mass.

To date, SSTR positivity in fibroadenoma is only shown in clinical cases. Here in this exploratory study, SSTR positivity is shown at both gene and protein levels in both fibroadenoma and phyllodes tumour tissues.

## MATERIALS AND METHODS

### Tissue collection and cell culture

Tissue samples were collected from patients who underwent tumorectomy and mastectomy for fibroadenoma and phyllodes tumour, respectively, at Acıbadem Mehmet Ali Aydınlar University Atakent Hospital and Koç University Hospital, Istanbul, Turkey. All patients were informed, and written consent was obtained. The studies were approved by the Ethics Committees of Acıbadem Mehmet Ali Aydınlar University (2022-15/17) and Koç University (2015.167.IRB2.064). Tumour samples were fresh-frozen in liquid nitrogen and stored at -80°C for protein isolation or preserved in RNAlater solution (Qiagen) for RNA isolation. Patients’ tissue slides were kindly provided by the Koç University Hospital Pathology Department.

Cell isolations for immunofluorescence experiments were performed using the explant (outgrowth) culture method as previously described^14^. Tumour tissues were cut as 1 mm x 1 mm dimension and cultured onto 1% gelatine coated slides and incubated at 37°C in a 5% CO_2_ incubator in an FBS enriched medium (DMEM High glucose supplemented with 20% foetal bovine serum and 1% penicillin/streptomycin, and 1% Amphotericin B). Tissues were cultured for 4-10 days.

### Immunofluorescence Analyses

Cells from the explant cultures were fixed with 4% paraformaldehyde, permeabilised with 0.1% Triton X-100, blocked with Superblock (ScyTEK) and incubated with anti-cytokeratin 18 (CK18) (1:50) (ab688, Abcam), anti-α smooth muscle actin (α-sma) (1:50) (M0851, Dako) or anti-SSTR2 (1:50) (ab134152, Abcam) primary antibodies for 90 minutes at 37°C. Following primary antibody incubations, cells were incubated with Cyanine3-labelled-goat anti-rabbit (Cy3) (1:100) (111-165-003, Jackson Laboratories), or fluorescein isothiocyanate–labelled-goat-anti-mouse (FITC) (1:100) (115-095-003, Jackson Laboratories) secondary antibodies for another 90 minutes at 37°C and mounted with 4’,6-diamidino-2-phenylindole (DAPI) containing mounting medium (Abcam) as published before^14^. Images are taken using the Leica DMI8/SP8 Confocal System.

### Immunohistochemistry (IHC) Analyses

Immunohistochemistry staining was performed as previously described with some modifications^15^. Briefly, sequential 3 μm thick paraffin-embedded tissue sections were placed on the same slide, de-paraffinized, and re-hydrated. Antigen retrieval was performed by boiling the slides in Antigen Retrieval Buffer (100x Citrate Buffer pH:6.0, Abcam) for 10 min in a microwave oven (600V). Mouse and Rabbit Specific HRP/DAB (ABC) Detection IHC kit (ab64264, Abcam) was used according to the manufacturer’s instructions for the staining. Primary antibodies used were anti-CK18 (1:1000) (ab688, Abcam), anti-α-sma (1:2500) (M0851, Dako), anti-SSTR2 (1:2500) (ab134152, Abcam), anti-CD31 (1:1000) (ab281583, Abcam), and anti-CD34 (1:1000) (MA1-10202, Invitrogen) along with Rabbit isotype control IgG (1:2500) Mouse isotype control IgG (1:2500). Tissue slides were incubated with primary antibodies overnight at 4°C. All washes were done with phosphate buffer supplemented with 0.1% Tween-20 (PBS-T). Following DAB substrate incubation, the counterstaining was performed using Mayer’s hematoxylin solution. A Zeiss Axiocam 3.1 system was used throughout the process.

### Western Blot Analyses

Fibroadenoma, phyllodes and PanNET tissues were first disintegrated using a blue pestle and then lysed in 1% sodium dodecyl sulphate (SDS) hot lysis buffer following the hot lysis protocol (Abcam). SSTR-positive PanNET tissue was used as a positive control. Protein concentration was determined by Bradford assay. 20 μg of protein extracts were mixed with 4x Laemmli sample buffer with 2-mercaptoethanol (Biorad) and heated at 95°C for 10 minutes. Proteins were separated using Any KD SDS-PAGE (Biorad) gel and transferred onto a PVDF membrane by the application of 250 mA for 90 minutes. The membrane was blocked with 5% non-fat milk dissolved in Tris-buffered Saline containing 0.1% Tween-20 (TBS-T) for 1 hour, then incubated with anti-SSTR (1:5000) (ab134152, Abcam) antibody overnight at 4°C, washed with TBS-T and incubated horseradish-peroxidase linked goat anti-mouse secondary antibody for 1 hour at room temperature. Antibody detection was performed using enhanced chemiluminescence; Supersignal West Femto Maximum Sensitivity substrate (Thermo Scientific). Equal loading was verified using an anti-GAPDH antibody (MA5-15738, Invitrogen) after mild stripping of the membrane. The chemiluminescence signal was detected using Chemidoc MP imaging systems (Biorad).

### RNA isolation, cDNA synthesis and Real-time polymerase chain reaction (RT-PCR)

mRNA extractions, preparations, and RT-PCR were performed as described previously^16^. Approximately 10 mg of tissues were used for RNA isolation. To isolate total RNA, the Zymo Quick-RNA Miniprep Kit was used, and the manufacturer’s instructions were followed. The amount and quality of the RNA were measured using a nanophotometer (Implen, Thermo Fischer). cDNA was synthesised using an M-MLV Reverse Transcriptase kit (Invitrogen) and 1000 ng RNA was used. The final volume was adjusted to 100 μl with nuclease-free water (Final cDNA concentration: 10 ng/ μl).

RT-PCR was carried out in Applied Biosystems Quantstudio 7 RT-PCR system (Applied Bioscience). Primer sequences used in this experiment were as follows: SSTR1 FWD: 5’-CACATTTCTCATGGGCTTCCT-3’ REV: 5’-ACAAACACCATCACCACCATC-3’ SSTR2 FWD: 5’-GGCATGTTTGACTTTGTGGTG-3’ REV: 5’-GTCTCATTCAGCCGGGATTT-3’ SSTR4 FWD: 5’-CGTGGTCGTCTTTGTGCTCT-3’ REV: 5’-AAGAATCGGCGGAAGTTGT-3’ SSTR5 FWD: 5’-CTGGTGTTTGCGGGATGTT-3’ REV: 5’-GAAGCTCTGGCGGAAGTTGT-3’. Total volume of 20 μl reaction mix contained 10 μl LightCycler^®^ 480 SYBR Green I Master Mix (Roche), 2 μl 2.5 mM primer (F+R), 2 μl cDNA (1000 ng/ 100 μl) and 6 μl nuclease-free water (Thermo Scientific). qRT-PCR cycles included pre-incubation at 95°C for 5 minutes followed by 40 cycles of 95°C for 10 seconds 60°C for 30 seconds and 72°C for 30 seconds. PCR products were also analysed by melting curves to confirm the specificity of the primers under reaction conditions. All melting curves revealed well-defined peaks with the expected melting temperatures. Controls that contain no cDNA were included in every run to monitor potential contamination. The threshold cycle number (Ct) represents the cycle number at which the amount of amplified target product reached a certain threshold.

All primers were designed to amplify the genes of interest and were optimised using cDNA isolated from RNA of SSTR-positive PanNET tissue. Linear regression was plotted using 1:1 serial dilution of the cDNA starting from 40 ng per 20 μl reaction mixture.

## RESULTS

### SSTR is expressed in both fibroadenoma and phyllodes tumours

RT-PCR and western blot were performed to assess SSTR expression in different tumour types (Figure 2). PanNET tissue, known to express SSTR2, was used as a positive control. Gene expression analyses revealed that fibroadenoma and phyllodes tumours exhibited positivity in all SSTR subtypes (Figure 2A). SSTR3 gene expression was not determined due to the primer’s low amplification efficiency. In fibroadenoma and phyllodes tissues, particularly SSTR4 and 5, were observed to be higher than in PanNET. HepG2, as a negative control, showed no expression of SSTR1-4, with minimal expression of SSTR5, confirming the information from The Human Protein Atlas regarding HepG2 cell lines (https://www.proteinatlas.org/ENSG00000162009-SSTR5/cell+line). SSTR2 protein was observed in both phyllodes tumour and fibroadenoma (Figure 2B).

**Figure 2:**
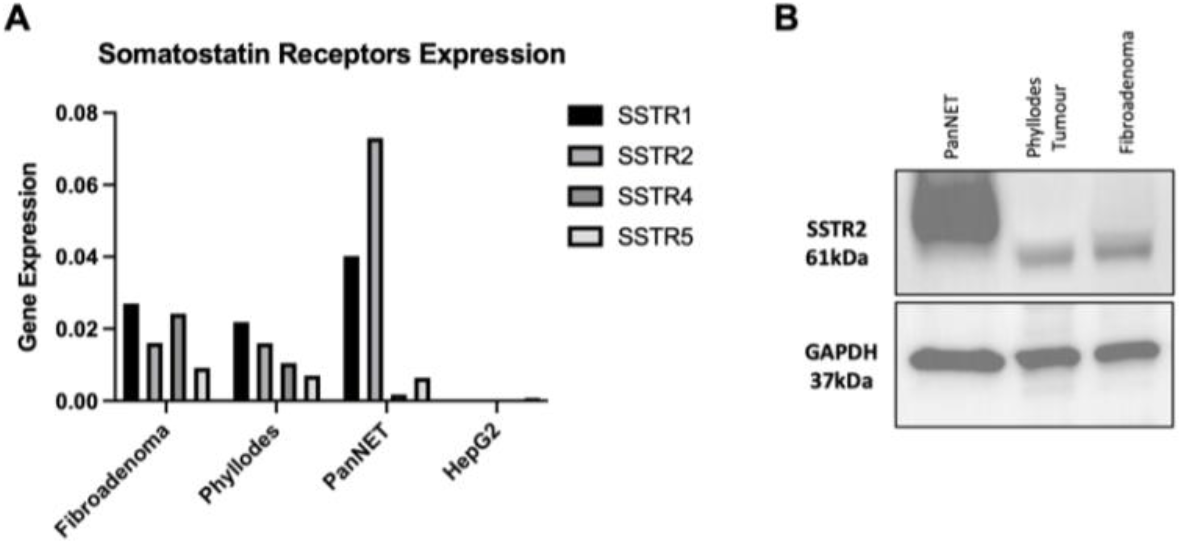
A) SSTR gene expression in tumour tissues of fibroadenoma, phyllodes and PanNET. HepG2 cell line was used as a negative control. B) SSTR2 protein expression in tumour tissues: PanNET, Phyllodes tumour and fibroadenoma. GAPDH was used for loading control.

### SSTR2 expression patterns in Fibroadenoma and Phyllodes tumours

Breast epithelium consists of distinct subpopulations: Luminal cells and basal/myoepithelial cells^17^. Luminal cells, identified by CK18 expression, are epithelial cells that line the ducts and are responsible for milk secretion^18^. Myoepithelial populations, situated between luminal cells and the stroma forming a monolayer over the ductal luminal epithelial cells, contribute to tissue integrity and communication between both compartments^18, 19^. Immunohistochemistry was performed to identify the cell population expressing SSTR2. Various markers including CK18 for luminal cells, α-sma for basal/myoepithelial cells, CD31 for endothelial cells, and CD34 for stromal fibroblasts were used to distinguish different cellular components in fibroadenoma and phyllodes tumours.

It is observed that unlike in neuroendocrine tumours (NET) (Supplementary Figure 1 shows SSTR2 expression in PanNET tumour as an example), SSTR2 was not expressed by the epithelial cells in fibroepithelial lesions (Figure 3A-D). The pattern of SSTR2 expression seemed to overlap with α-sma-positive areas; therefore, attention was focused on the areas predominantly expressing α-sma, such as myoepithelial cells around the luminal cells. As depicted in Figure 3, CK18-positive luminal cells (A) were surrounded by α-sma-positive myoepithelial cells (B); however, SSTR2 expression was not detected in these cells (D). Other areas that were α-sma-positive were blood vessels and capillary structures. They were identified by the inner layer of endothelial cells positive for CD31, and the outer layer of pericytes positive for α-sma. Figure 3E-F shows a blood vessel structure. As it depicted SSTR2 expression seemed to overlap within these two layers as part of the blood vessel structure. Stromal fibroblasts did not seem to express SSTR2 as no specific overlap was observed in CD34-positive areas, except the parts that stained hematopoietic cells, which overlap with CD31.

**Figure 3:**
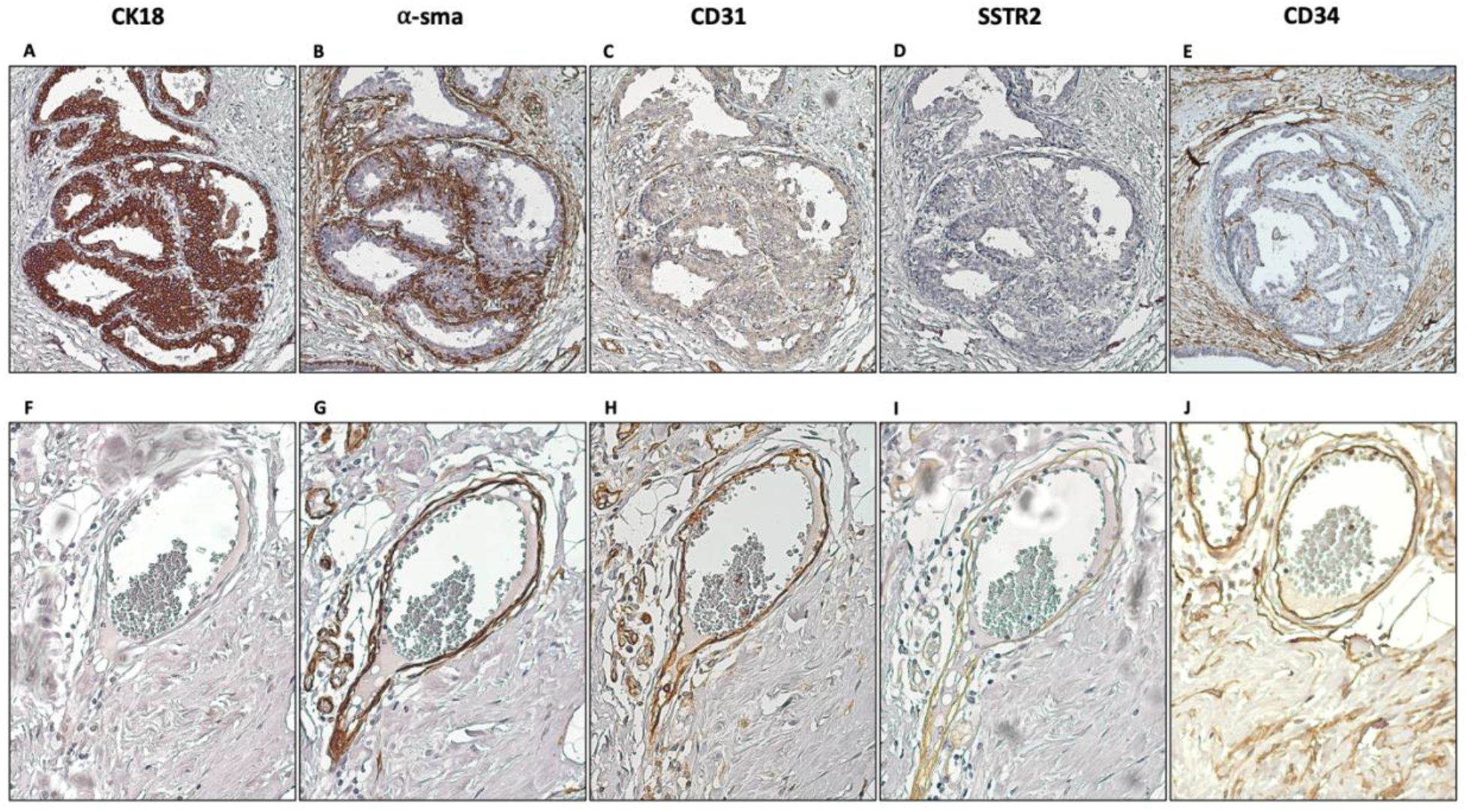
Cellular component that expresses SSTR2. CK18 (A, E), α-sma (B, F), CD31 (C, G), SSTR2 (D, H), and CD34 (E, J) were stained in consecutive tissue slides and staining patterns were investigated. Neither luminal cells (characterised by CK18 expression (A), nor the adjacent myoepithelial cells which express α-sma (B) showed SSTR2 expression (D). On the contrary image E-H shows a blood vessel that contains endothelial cells expressing CD31 (G), and adjacent pericytes expressing α-sma (F) show SSTR2 positivity. CD34 staining shows the stromal fibroblasts which also lack SSTR2 expression. (Magnification 20x)

As the next step, we wanted to see if this pattern of expression is the same in all fibroepithelial lesions. Because the phyllodes tumours are rare, we were able to analyse a borderline and a malignant phyllodes subtype along with a fibroadenoma. It is observed that the SSTR2 expression pattern did not differ among tumour types, which was confined to blood vessel and capillary structures (Figure 4).

**Figure 4:**
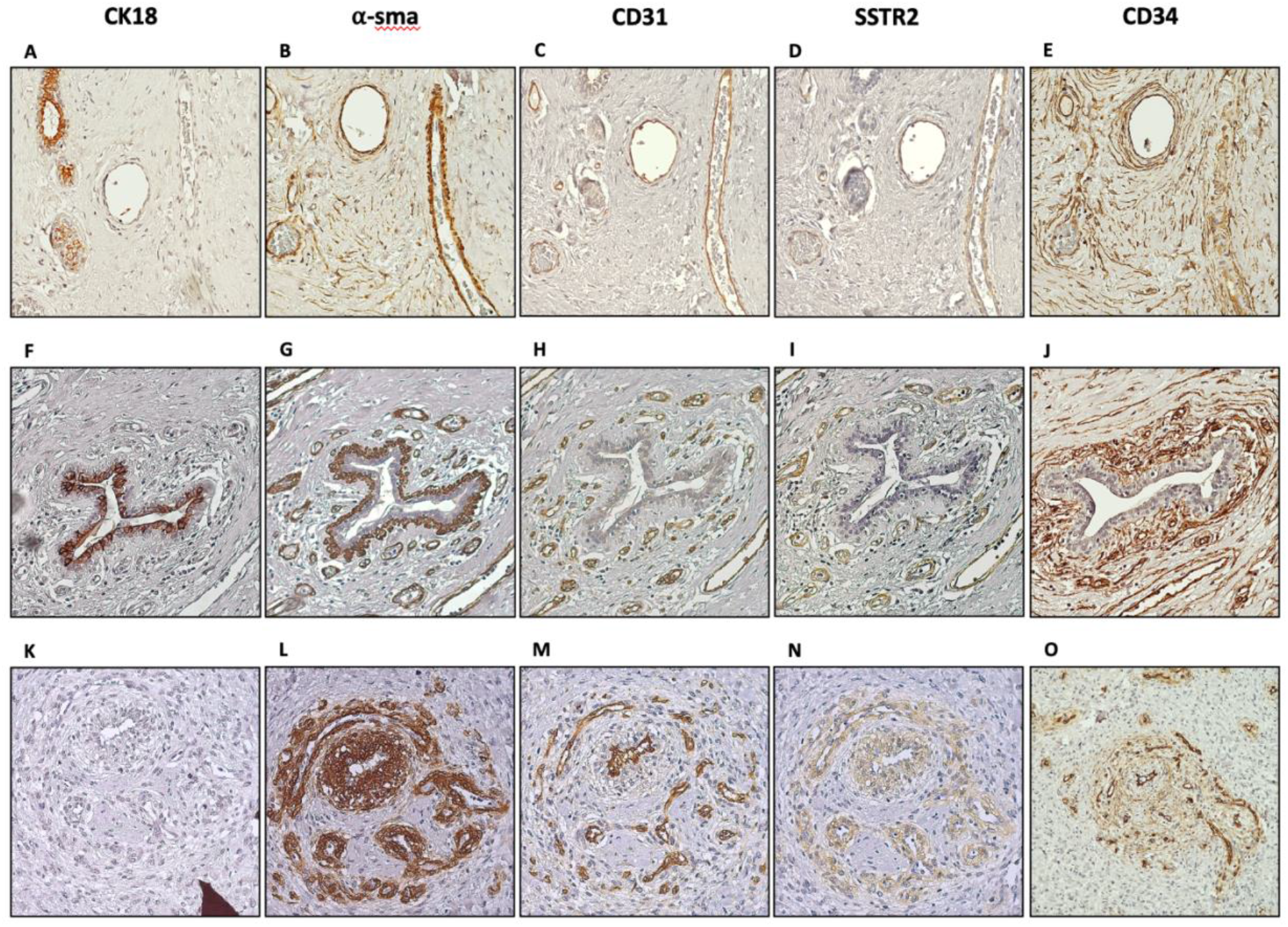
Characterisation of SSTR2 expressing group of cells in fibroadenoma (A-E) and phyllodes tumours: Borderline (F-J) and Malignant (K-O). CK18 (A, F, K), CD34 (B, G, L), α-sma (C, H, M), CD31 (D, I, N) and SSTR2 (E, J, O) were stained in consecutive tissue slides and staining patterns were investigated. (Magnification 10x)

For further characterisation, a primary explant culture was performed to obtain cells from the fibroadenoma tissue. The growth medium was not selected to favour a specific type of cells to obtain a mixed-cell culture. Supplementary Figure 2 shows the progression of the outgrowth culture by days and how the mixed-cell culture was formed. The explant culture was terminated on day 7-10 right before cobblestone/round-shaped cells were starting to detach from the plastic surface. Afterwards, cells were fixed with 4% paraformaldehyde, and immunofluorescence staining was performed. While α-sma-positive cells did not show SSTR positivity, cells with cobblestone morphology with large nuclei with faint CK18 expression showed pronounced SSTR2 expression on their surfaces (Figure 5). Even though the phenotypes of these cells are strong indicators of endothelial cells, to conclude, CD31 staining needs to be performed.

**Figure 5:**
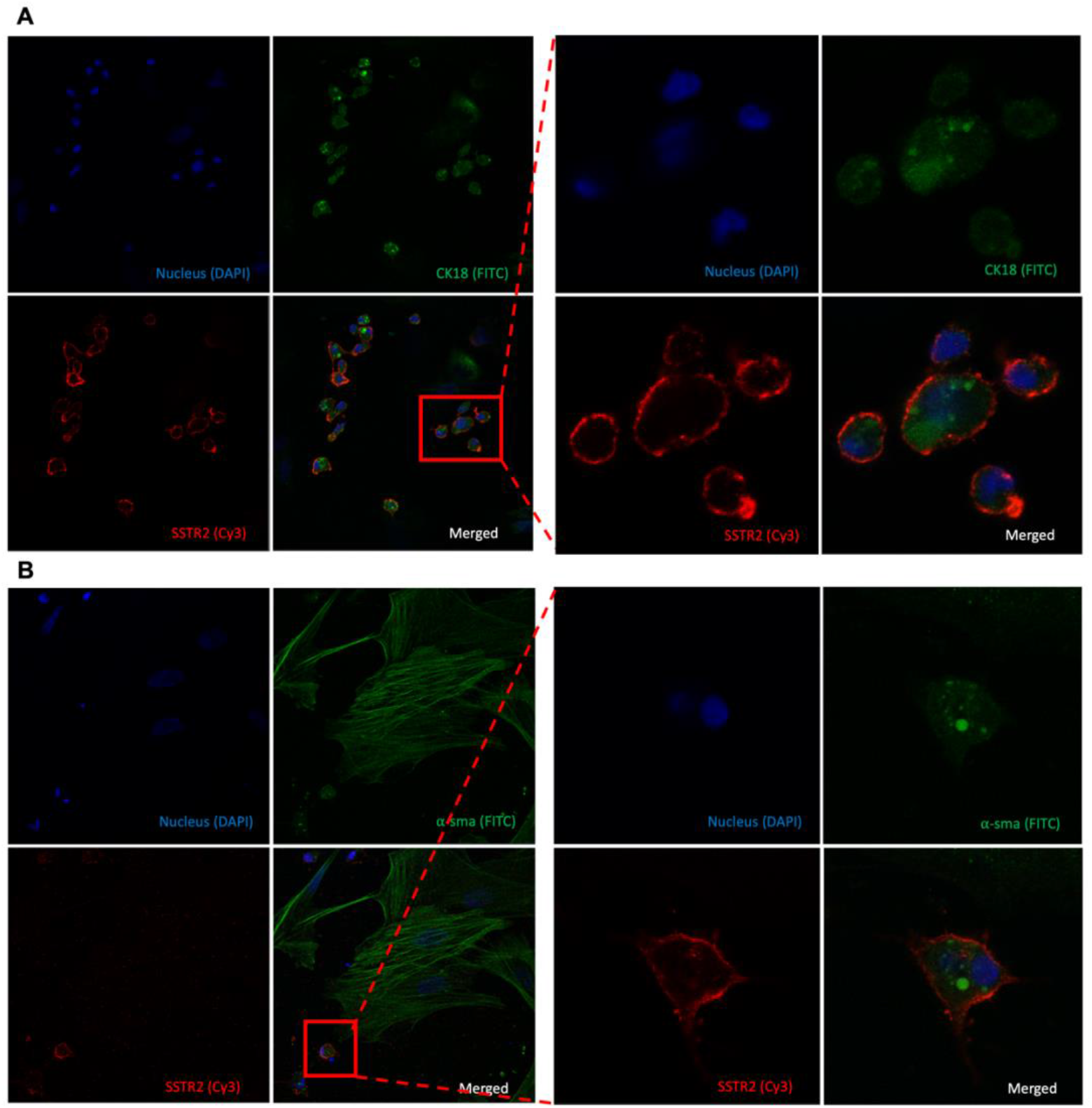
Characterisation of SSTR2 expressing group of cells in primary fibroadenoma mixed culture. Double staining of CK18 (FITC, green) and SSTR2 (Cy3, red) (A) with 40x magnification on the right. Double-staining α-sma ((FITC, green) and SSTR2 (Cy3, red) (B) with 40x magnification on the right.

## DISCUSSION

Fibroadenoma and phyllodes tumours are fibroepithelial lesions of the breast, representing a spectrum of lesions from benign with minimal recurrence potential to relatively aggressive examples (“cystosarcoma”) with progression and metastasis. Both are characterised by biphasic patterns composed of stromal and epithelial cells. While fibroadenomas are common benign tumours, phyllodes tumours are rare, constituting 0.3-1% of all breast tumours and representing a spectrum of lesions with varying clinical behaviour. Fibroadenoma can even be managed with a wait-and-watch approach without surgery, including follow-ups, vacuum-assisted biopsy and cryoablation, preserving the breast and maintaining patients’ quality of life. On the other hand, for phyllodes tumours, regardless of subtypes, the only treatment is surgical excision with a tumour-free margin, often resulting in complete mastectomy^13^.

SSTR-expressing tumours have been successfully diagnosed and treated using DOTATATE radiotracer systems, particularly in patients with NETs. These theranostic systems are highly specific to tumours with high SSTR expressions. Although SSTR expression is observed in breast adenocarcinomas (epithelial malignancies of this organ), when it comes to fibroepithelial lesions, only two incidental cases have been reported in the literature thus far, both fibroadenomas, and both were detected during NET screening. A similar case, in which the patient’s fibroadenoma shrank during ^177^Lu treatment for PanNET, prompted us to conduct this more detailed analysis of fibroepithelial tumours to investigate the potential use of this incidental finding for diagnostic and therapeutic purposes.

Accordingly, we have investigated SSTR expression in fibroepithelial lesions at a cellular and molecular level. This is the first in the literature to reveal that these receptors are indeed expressed in phyllodes tumours, illustrating that it is the stomal cells of these fibroepithelial lesions, not the epithelial component, that corresponds to DOTATATE positivity at the clinical/imaging level. We observed that SSTR in fibroepithelial lesions, regardless of the aetiology, may be found around the blood vessel and capillary structures and seem to overlap with CD31-positive endothelial cells and adjacent α-sma positive pericytes. It is rather difficult to ascertain which cell types in the stroma are specifically expressing the SSTR with the information we gathered from our experiments, however the expression was in the region where stromal cells are located, and the expression was negative in the epithelial component.

Of note, IF staining illustrated that SSTR was present in the cells that have endothelial cell morphology as well. Whether these vascular elements are an integral part of the “stromal” component of the fibroepithelial lesion or whether they are the native vessels, it is conceivable that differential tendency may be a factor. Supporting this preferential expression in the possible tumour-region endothelial cells, Kumar et al. also found SSTR2 expression in the blood vessels and surrounding smooth muscle cells, also known as pericytes, in human breast cancer alongside tumour cells^20^. In the literature, endothelial cells are shown by many studies to express SSTR2 in diseases, and most of those studies associated SSTR2 expression in endothelial cells with angiogenesis. Interestingly, studies that evaluate endothelial cells in general find different SSTR subtypes more dominant, such as SSTR1, SSTR4 and SSTR5^21, 22^, while other studies that investigate proliferative situations, such as inflammatory conditions or wound healing, find SSTR2 predominantly expressed in endothelial cells^23^. In support of that, SSTR2 and 5 are expressed in angiogenesis during the proliferation of endothelial cells^24^. Similarly, Watson et al. revealed that only growing endothelial cells express SSTR2; therefore they suggested that since an angiogenic response is required for tumours to grow more than 2 mm in diameter, SSTR2 antagonists could be a potential therapeutic approach for tumours larger than 2 mm^25^. It is well-documented that somatostatin or its analogues such as octreotide, have anti-proliferation activity in endothelial cells^26, 27^ Therefore, in our case, ^177^Lu-DOTATATE might have acted as an anti-angiogenic treatment, potentially resulting in tumour shrinkage, as disrupting the blood supplies or preventing the formation of new blood vessels can be an efficient approach to reducing tumour sizes.

While many studies associated SSTR2 expression with endothelial cells, only recent studies have found pericytes to be associated with SSTR2; for instance, SSTR2 has been related to active inflammation in temporal arteritis and was mainly detected in macrophages and pericytes. Additionally, a single-cell proteomics study showed that pericytes are the most strongly associated cells with SSTR2 in lung fibrosis in inflammatory lung diseases^28^. Pericytes play an essential role in angiogenesis, such as expressing vascular endothelial growth factor, guiding vessel sprouts by migrating ahead of endothelial cells, and vessel maturation in the context of neovascularisation^29^. Even though so far there is no direct correlation between SSTR2 expression and pericytes in cancer, pericytes help maintain the integrity and functionality in tumour vasculature; therefore depleting pericytes make blood vessels within the tumour more vulnerable and sensitive to therapies^29-31^.

These findings shed new light to the progression of fibroepithelial neoplasms. It has long been a curiosity as to which component of these tumours is truly involved in the driver’s seat. The fact that malignancy in these tumours typically arises from the stromal component and manifests as sarcoma has suggested that, in fact, it may be the stromal component that is the main driver in these tumours. The preferential expression of SSTRs in the stromal component also supports this impression.

In this study, we confirmed the SSTR2 expression pattern in fibroadenoma, borderline and malignant subtypes of phyllodes tumours. However, further investigation is needed to validate this expression pattern in more samples, including benign subtypes of phyllodes. This study stands out by revealing the existence of SSTRs in challenging phyllodes tumours and demonstrating that the receptors are located in the same area as in fibroadenoma. Given our observation of tumour shrinkage in fibroadenoma during ^177^Lu treatment, there might be a possibility that these receptors in phyllodes tumours are also functional, similar to those in fibroadenoma. Therefore, this study may have uncovered new treatment opportunities for patients with these neoplasms without hampering their quality of life.

## CONCLUSION

Herein we examined SSTR expression at both gene and protein levels in both fibroadenoma and phyllodes tumours. While previous research has exclusively demonstrated SSTR expression in fibroadenoma through DOTATATE tracer positivity in clinical studies and case reports, our study is the first to report SSTR expression in phyllodes tumours in the existing literature.

Another important finding of the current study is that ^177^Lutetium treatment causes shrinkage not only in the aimed target of neuroendocrine metastasis but also, as an off-target, in breast fibroadenoma. This discovery suggests a potential avenue for exploring non-surgical treatment options for these rare cancers. However, it is crucial to note that our study, while promising, was exploratory in nature and limited by sample size. Further comprehensive evaluations are needed to validate and expand upon our findings.

## DECLARATIONS

### Ethics Approval and Consent

This study was approved by the Ethical Committee of Acıbadem Mehmet Ali Aydınlar University (2022-15/17) and Koç University (2015.167.IRB2.064) and performed following the Declaration of Helsinki.

All patients were given written informed consent. A study nurse accompanied all the patients if they needed further questions about the consent forms. By the end of this process, patients who agreed to participate in the study signed a consent form. The ^177^Lu-DOTATATE image that is used in the manuscript was included after the written consent was obtained from the patient. All patients’ details were anonymised upon participation in the study.

## Acknowledgement

We would like to express our sincere gratitude to Prof. Nazmi Volkan Adsay from the Department of Pathology, Koç University School of Medicine for his valuable discussion, scientific contribution and material support. We would also like to thank our former research group members Orhan Ağcaoğlu and Setenay Gupse Özcan for their assistance in the experimental parts of the study.

## Availability of Data and Materials

All data generated or analysed during this study are included in this published article.

## Authors’ Contribution

HS: Investigation, Methodology, Writing-Original Draft, Visualisation, Review and Editing

ME: Conceptualisation, Resources, Supervision, Writing-Review and Editing

## Supplementary Figures

**Supplementary Figure 1:**
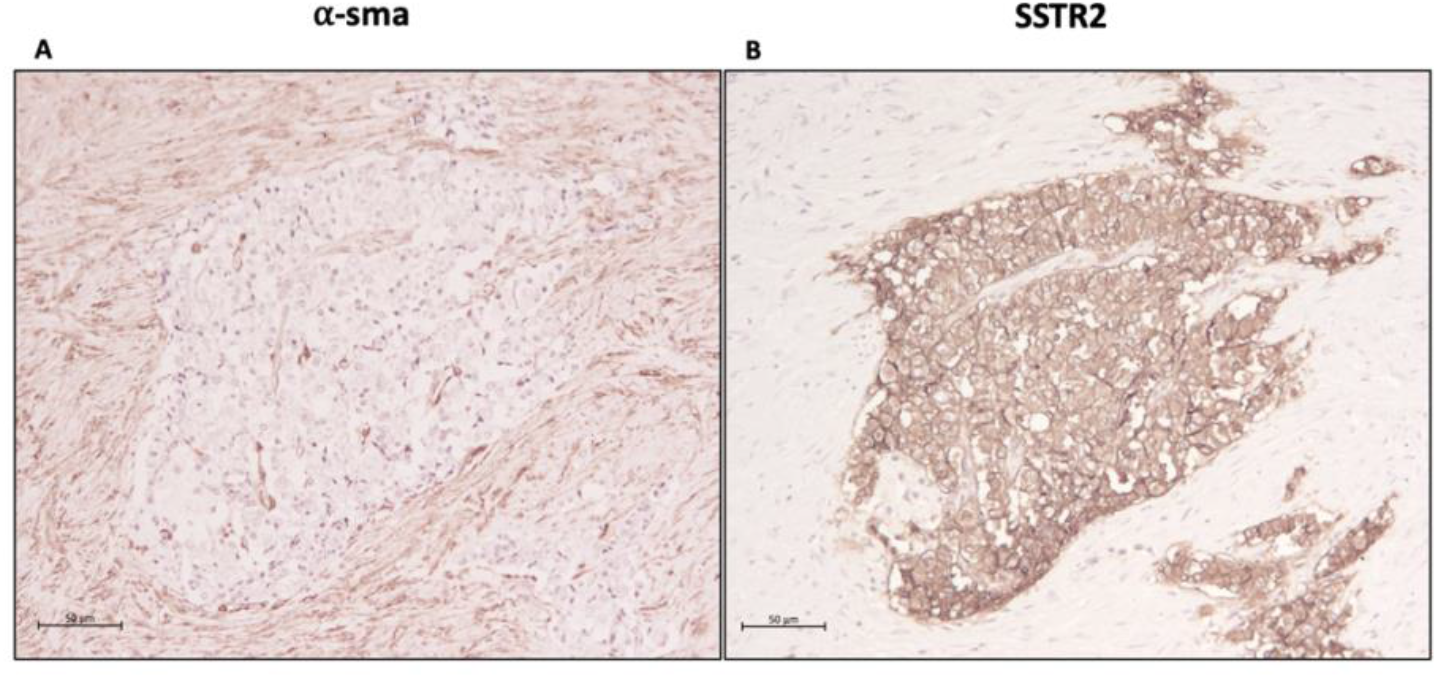
In PanNET tumours SSTR2 expression is confined to epithelial cells. α-sma-positive cells do not express SSTR2.

**Supplementary Figure 2:**
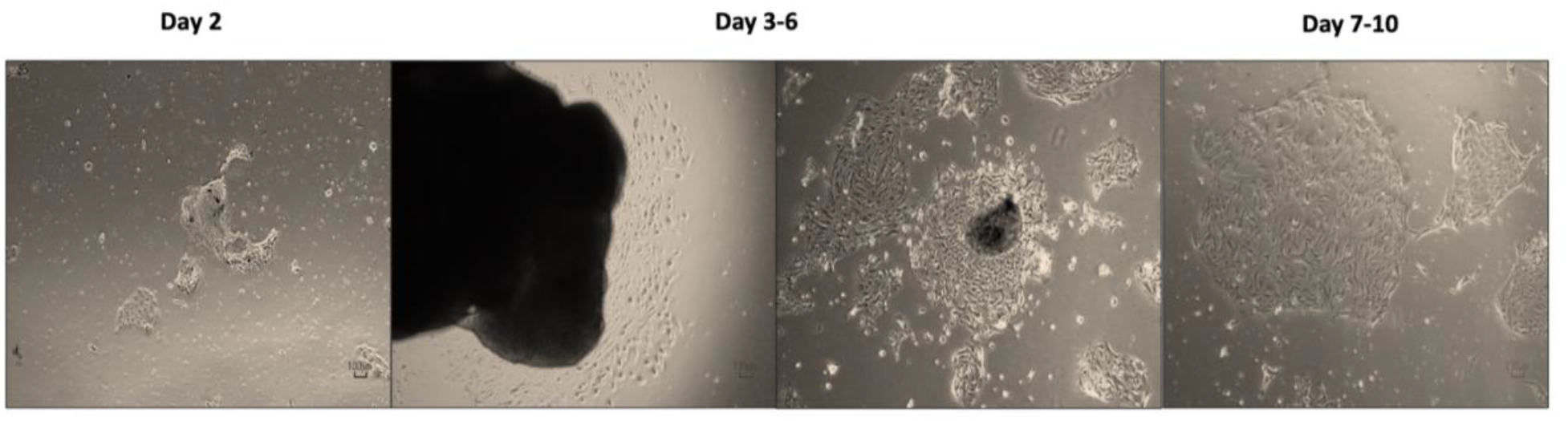
Cells outgrown from the explants constituting a heterogenous cell population containing epithelial-like cobblestone-shaped cells and spindle-shaped mesenchymal cells.

